# Rapid, multiplexed, whole genome and plasmid sequencing of foodborne pathogens using long-read nanopore technology

**DOI:** 10.1101/558718

**Authors:** Tonya L. Taylor, Jeremy D. Volkening, Eric DeJesus, Mustafa Simmons, Kiril M. Dimitrov, Glenn E. Tillman, David L. Suarez, Claudio L. Afonso

**Author notes:** Corresponding author: **Claudio L. Afonso**. Mailing address: USDA ARS, Southeast Poultry Research Laboratory, 934 College Station Rd., Athens, GA 30605.

## Abstract

United States public health agencies are focusing on next-generation sequencing (NGS) to quickly identify and characterize foodborne pathogens. Here, the MinION nanopore, long-read sequencer was used to simultaneously sequence the entire chromosome and plasmids of *Salmonella enterica subsp. enterica* serovar Bareilly and *Escherichia coli* O157:H7. A rapid, random sequencing approach, coupled with *de novo* genome assembly within a customized data analysis workflow, that can resolve highly-repetitive genomic regions, was developed. In sequencing runs, as short as four hours, using nanopore data alone, full-length genomes were obtained with an average identity of 99.87% for *Salmonella* Bareilly and 99.89% for *E. coli* in comparison to the respective MiSeq references. These long-read assemblies provided information on serotype, virulence factors, and antimicrobial resistance genes. Using a custom-developed, SNP-selection workflow, the potential of the nanopore-only assemblies (after only 30 minutes of sequencing) for rapid phylogenetic inference, with identical topology compared to the published dataset, was demonstrated. To achieve maximum quality assemblies, the developed bioinformatics workflow employed additional polishing steps to correct the systematic errors produced by the nanopore-only assemblies. Nanopore sequencing provided a shorter (10 hours library preparation and sequencing) turnaround time compared to other NGS technologies.

## Introduction

United States public health agencies routinely perform surveillance on microbial foodborne pathogens, and in the U.S. alone each year, approximately 1 in 6 individuals are sickened by foodborne illnesses, resulting in approximately 3,000 deaths^1^. During outbreak responses, identification of the source is instrumental for the fast removal of the contaminated items from public circulation. However, specific characterization of foodborne pathogens during these surveillance programs in food production and distribution is important, as it allows for early warnings and possible removal of the contaminated food product(s) before the development of an outbreak^1^.

To that end, U.S. public health agencies have employed next-generation sequencing (NGS) using short-read sequencing technology in surveillance activities and outbreak response^2^. In addition to utilizing whole genome sequencing (WGS) for pathogen identification, more detailed information on the pathogen such as virulence, antimicrobial resistance, serotype, and inference of possible links between the sources of contamination is obtained^3^. WGS has provided faster identification of pathogens from contaminated sources of outbreaks, reduced the number of illnesses and deaths due to the foodborne infections, and decreased the number of isolates needed to link the illness to the source of contamination^4,5^.

Although WGS is now a routine procedure in epidemiologic investigation and surveillance of foodborne pathogens, short-read, sequencing technology present challenges such as resolving repetitive regions, leading to incomplete *de novo* assemblies (severe fragmentation)^6-8^. These gaps can lead to the inability to determine genome organization or architecture, which can be important in determining if genes are co-regulated or co-transmissible in the case of genes associated with mobile elements^9^. Even though the short-reads are accurate, closed whole genome assemblies are now commonly accomplished using a combination of both short-read (for base accuracy) and long-read sequencing technologies (for structural accuracy)^10–12^.

Long-read sequencing, enabled by single-molecule real-time (SMRT) sequencing technology that has been utilized since 2004, can produce reads averaging 11kb in length, which facilitates the completion of bacterial genome assemblies that are either lacking in sequencing depth at certain repetitive areas of the genome or have areas that are missing reads completely using short-read technology^13^. The long-reads span across these large repetitive regions^14–16^ and can provide unbiased coverage of regions sequenced poorly with other technologies due to G/C content or other characteristics^13,17^. However, there is still a need for an approach that generates inexpensive, long-read data in a short turnaround time to be utilized for both rapid detection of an organism, complete sequencing of bacterial chromosomes and plasmids, and complementation to other sequencing technologies used in both outbreak investigations and foodborne pathogen surveillance.

Oxford Nanopore has developed technology to fit this role in the form of the MinION nanopore-based sequencer that produces long, single-molecule reads. This novel sequencer is pocket-size (10cm x 2cm x 3.3cm) and powered directly by a USB port from a laptop computer^18^. It is portable, field-deployable, inexpensive, and can provide sequencing of both DNA and RNA in real time. Nanopore sequencing still produces systematic errors, and for that reason, it has previously only been used as a complement of short read sequencing. The use of nanopore-only assemblies for full-length genome sequencing, genome structure determination, antimicrobial resistance gene identification, and phylogenetic analysis would be a significant advance that would allow the universal low-cost access of information necessary for critical management decisions. Since the release of the MinION platform, bioinformatic tools have been steadily evolving, with the goal of using nanopore data to assemble accurate, whole, bacterial genomes independent of any other sequencing technology^19^.

In this study, utilizing the nanopore technology, we aimed to simultaneously sequence and assemble complete genomes of two pathogenic bacterial strains that can cause human illness worldwide, *Salmonella enterica subsp. enterica* serovar Bareilly and *Escherichia coli* O157:H7. Using a custom, reproducible bioinformatics workflow that employs publicly-available tools, the circularized bacterial genomes and associated plasmids of both strains were assembled and polished with a final error rate of only 0.1 %. This study shows that long-read nanopore sequencing can be used as a low-cost method to generate closed assemblies of microbial foodborne pathogen genomes and associated plasmids and demonstrates that the data is of sufficient quality for phylogenetic classification of *Salmonella* isolates. These closed assemblies provide information on genome organization and can complement existing characterization data from other technologies such as short-read sequencing.

## Materials and Methods

### Bacterial cultures and DNA extraction

The *Salmonella* Bareilly isolate (CFSAN000189) was isolated from raw shrimp in India (Biosample SAMN04364135), and the *E.coli* O157:H7 isolate (FSIS11705876) was isolated from domestic, raw, ground beef collected by the U.S. Department of Agriculture Food Safety and Inspection Services (USDA-FSIS) as part of routine sampling of a U.S. establishment (Biosample SAMN08167607). Both bacterial isolates were grown on sheep blood agar (SBA) for 24 hours at 35 °C. Total DNA from each isolate was extracted using the DNeasy Blood and Tissue Kit (Qiagen, USA) following manufacturer’s instructions. DNA concentrations throughout the experiment were determined by using the Qubit^®^ dsDNA HS Assay Kit on a Qubit^®^ fluorometer 3.0 (Thermo Fisher Scientific, USA).

### Library Preparation and MinION Sequencing

The 1D gDNA long read selection protocol was used with the SQK-LSK108 kit (Oxford Nanopore Technologies (ONT), UK) to prepare MinION-compatible libraries. The DNA shearing step was eliminated from the protocol with the aim of selecting for very long reads. Approximately, 2μg of *E. coli* DNA and 2 μg of *Salmonella* DNA in a total of 100 μL each were added to the NEBNext^®^ Ultra™ II End Repair/dA-Tailing module (New England Biolabs (NEB), USA) for end repair and dA-Tailing, following manufacturer’s instructions, and purified using Agencourt AMPure XP beads (Beckman Coulter, USA). Each purified, end-prepped DNA product was barcoded using a separate barcode from the 1D Native barcoding kit (EXP-NBD103, ONT, UK) and following the 1D Native barcoding genomic DNA protocol. The samples were then bead-purified (Beckman Coulter, USA), and equimolar amounts of each barcoded sample were pooled together for a final quantity of 700ng. Adapters were ligated to the pooled sample using Blunt/TA ligase (NEB, USA) following the 1D gDNA long read selection protocol. The MinION device was used to sequence the created library on a new FLO-MIN106 R9.4 flow cell^20,21^. The standard 48 hr 1D sequencing protocol was initiated using the MinKNOW software (ONT, UK). Average quality and coverage of the raw sequencing data were determined using CG-pipeline^22^.

### MiSeq Sequencing and Quality control

To verify the newly developed approach used in this study, libraries for short-read WGS of the *Salmonella* Bareilly and *E. coli* isolates were prepared using the Nextera XT kit (Illumina, USA) according to the manufacturer’s protocol. The libraries were loaded separately into a single flow cell of the 300 and 500 cycle MiSeq Reagent Kits v2 for *Salmonella* Bareilly and *E. coli*, respectively, and paired-end sequencing (2×150 bp for *Salmonella* Bareilly and 2×250 bp for *E. coli)* was performed on the MiSeq instrument (Illumina, USA). The produced raw data were analyzed using SPAdes version 3.71^23^. Average quality and coverage of the raw sequencing data were determined using CG-pipeline^22^.

### MinION Bacterial Bioinformatics Workflow for Whole Genome Assembly

To analyze the MinION sequencing data, a customized workflow was developed. For subsequent time analysis, the data was also analyzed at intervals from the start of the sequencing. Reads were basecalled using Albacore (ONT v 2.0.2b) and subsampled for assembly using Filtlong (v.0.2.0)^24^ to a target depth of 75X. Fitlong subsampling is not random but keeps the longest and highest quality reads from the input, which targets maximum sequencing depth (total bases). Read quality was weighted more heavily than length (‘mean_q_weight 5), as testing showed this was necessary to retain sufficient coverage of small plasmids. The filtered reads were assembled using the Unicycler pipeline (v.0.4.7)^25^. This pipeline utilizes a minimap2/miniasm/racon iterative approach to assemble long-read-only data. Since Unicycler sometimes fails to detect valid end overlaps, assemblies were circularized using a custom script based on minimus2 (available in the workflow source repository)^26^. Circular contigs were rotated to start at a fixed position based on the reference. The consensus sequences were subjected to two rounds of polishing using Nanopolish (v.0.10.2)^27^, for which the full run (subject to time-based sub-setting but prior to FitLong subsampling) was used, and Benchmarking Universal Single-Copy Orthologs (BUSCO v.3.0.2)^28^ was used to evaluate the completeness of coding sequences and degree of gene fragmentation in the polished assemblies. To evaluate assembly accuracy, two procedures were used. For the *Salmonella* Bareilly isolate, which has previously been sequenced and published^29^, DNAdiff (MUMMER v.3.23)^30^ was used to evaluate both base-level and structural accuracy in the MinION assembly compared to the published reference. For the *E.coli* isolate, lacking a published reference, Illumina MiSeq reads were mapped to the assembly using BWA (v0.7.17), and LoFreq (v.2.1.3.1)^31^ was used to call single nucleotide polymorphisms (SNPs) and small indels, from which the assembly accuracy was calculated. Utilizing the short-read data, Pilon (v1.2.2)^32^ was used to error-correct small errors (‘--fix bases’) in the assemblies using existing short-read data from the same isolates prior to GenBank submission (SRA accession SRR498276 for *Salmonella* Bareilly; SRA accession SRR6373397 for *E. coli* O157:H7).

### MinION Annotation

The polished-MinION assemblies after 4 hours of sequencing were initially annotated using the “Annotate From” tool within Geneious 11.1.5 and the published *Salmonella* Bareilly strain CFSAN000189 (GenBank Accession NC_021844) and *E. coli* O157:H7 strain 9234 (GenBank Accession CP017446) sequences as references. ResFinder v.3.1 was used to locate any antimicrobial resistance genes and any point mutations that would result in antimicrobial resistance^33^. Additionally, to confirm the 4-hour assembly annotation, the pilon-corrected, final genome sequences were submitted to GenBank to be processed through the NCBI Prokaryotic Genomic Annotation Pipeline (PGAP) before being released.

### Phylogenetic Analysis

Twenty-three *Salmonella* reference datasets were downloaded (Supplementary Table 4) and used in tracing a foodborne outbreak in the U.S that were previously published^29,34^. The eight sub-sampled (15 mins to 1500 mins) unpolished S. Bareilly assemblies from this experiment were used to generate simulated Illumina datasets using ART (150 × 2, 50X coverage, MiSeq platform, 300 bp mean fragment length, 50 bp standard deviation)^35^. All datasets were analyzed with a SNP-calling pipeline using strain CFSAN000212 as a reference. Briefly, reads were optionally trimmed using Trim Galore (Illumina datasets), aligned to the reference using BWA-MEM^36^, called using LoFreq^31^, and filtered using local scripts according to specific criteria. For Illumina datasets, the VCF files were filtered by removing indels as well as any SNP with an alternate allele frequency of < 90%. Sites meeting one or more of the following criteria were flagged as suspect, and these loci were ignored during matrix generation: i) sites within 3 bp of a homopolymeric stretch of 4 bp or more; ii) sites occurring in a variant cluster (multiple variants within 2 bp of each other; iii) sites within 10 bp of a dam or dcm methylation motif; and iv) sites with observed A->G or T->C transition mutations. The remaining SNPs (64) were used to create a matrix of variable sites for phylogenetic reconstruction. PhyML v3.0 was used to generate a maximum likelihood SNP tree using the HYK85 model with 500 bootstrap iterations^37^. In this analysis, the Illumina result from the CFSAN000189 strain was replaced with the MinION-only assembly from sequencing the same strain in this study to ensure that our data would not be topologically attracted by the reference sequence and would cluster correctly without it. This tree was compared against the standard reference tree (utilizing the 23 reference strains) provided by Timme et al., 2017 ^34^.

### Availability of workflows, tools and code

The full NextFlow workflow, Conda environment configuration, and other associated code used in the analyses are publicly-available on GitHub (https://github.com/jvolkening/minion_bacterial).

## Results

### Analysis of MinION and MiSeq Raw Data

Before subsampling of the reads, the raw MinION sequencing data was used to estimate the mean depth for *Salmonella* Bareilly and *E. coli*, respectively. A total of 2.8 billion bases from 333,298 *Salmonella* Bareilly reads, with an average read length of 8638 nucleotides (nt), yielded a mean depth of 599X. For *E. coli*, a total of 3.8 billion bases from 429,909 reads with an average read length of 8979 nt were sequenced, and the mean depth was calculated to be 692X (Table 1). The shortest MinION read was 85 nt, which was from the *E. coli* isolate, while the longest read was from *Salmonella* Bareilly and was 129,119 nt. Both sets of MinION data had a mean read quality score above the standard (Q≥10), which indicates superior quality.

**Table 1:** Comparison of the Final Raw Data from MinION and Illumina

| Sequence Method | Average Read Length | Total Bases | Min Read Length | Max Read Length | Average Read Quality | Read Number | Mean Depth |
| --- | --- | --- | --- | --- | --- | --- | --- |
| MiSeq (*Salmonella*) | 149.51 | 288,633,579 | 35 | 151 | 36.66 | 1,930,511 | 57.72 |
| MinION (*Salmonella*) | 8638.36 | 2,879,148,408 | 113 | 120,119 | 19.36 | 333,298 | 599.06 |
| MiSeq (*E.coli*) | 242.61 | 556,035,081 | 35 | 251 | 34.96 | 2,291,825 | 111.2 |
| MinION (*E.coli*) | 8979.55 | 3,860,389,678 | 85 | 112,643 | 19.38 | 429,909 | 692.19 |

a. MiSeq Quality Standards = Q<sub>≥</sub>30
b. MinION Quality Standards = Q<sub>≥</sub>10

Illumina MiSeq data was also analyzed using the same bioinformatic tool. The MiSeq raw data had a depth of 57X for *Salmonella* Bareilly and 111X for *E. coli*. This sequencing technology produced 288 million bases from 1,930,511 *Salmonella* Bareilly reads, with an average read length of 150 nt. For *E. coli*, a total of 556 million bases from 2,291,825 reads were sequenced that had an average read length of 243 nt (Table 1). The minimum read length from both sets of bacterial sequences was 35 nt, while the longest was 151 nt for *Salmonella* Bareilly and 251 nt for *E. coli;* the MiSeq mean read quality was above the Q30 benchmark, signaling exceptional data.

### Assembly of MinION sequencing data

The raw MinION data for both isolates were subsampled on the basis of cumulative run time in order to simulate the effect of run length on final assembly quality. Subsets of reads generated in the first 15, 30, 60, 120, 240, 480, and 960 minutes (mins), in addition to the full run length, were analyzed (Tables 2 and 3). Four hours (240 mins) was determined as the shortest run time sufficient to assemble circular sequences from all chromosomes and plasmids from both isolates and represented a point after which longer run times resulted in remarkably diminishing gains in final accuracy (Fig S1). Data at each of the other run time subsets is available in Tables 2, 3, S2 and S3; however, the following analyses herein refer to the data collected in the first 240 mins of sequencing.

**Table 2:** Assembly Data for MinION sequencing of *Salmonella*

| Duration (min) | Reads | Subsampled Reads | Assembly Size | Circular Contigs<sup>a</sup> | Linear Contigs | Longest Contig | Longest Circular Contig | NG50<sup>b</sup> | Average identity in % | Reference Coverage |
| --- | --- | --- | --- | --- | --- | --- | --- | --- | --- | --- |
| 15 | 7229 | 7229 | 1135723 | 0 | 19 | 163786 | 0 | 0 | 99.13 | 24.36 |
| 30 | 14888 | 14888 | 4577215 | 0 | 18 | 841969 | 0 | 471499 | 99.55 | 95.38 |
| 60 | 29132 | 29132 | 4722179 | 1 | 0 | 4722179 | 4722179 | 4722179 | 99.79 | 98.4 |
| 120 | 51226 | 51226 | 4805334 | 2 | 0 | 4723663 | 4723663 | 4723663 | 99.84 | 100 |
| 240 | 84156 | 28492 | 4806150 | 2 | 0 | 4724389 | 4724389 | 4724389 | 99.87 | 100 |
| 480 | 132137 | 20193 | 4806518 | 2 | 0 | 4724724 | 4724724 | 4724724 | 99.87 | 100 |
| 960 | 248910 | 16221 | 4806892 | 2 | 0 | 4725103 | 4725103 | 4725103 | 99.89 | 100 |
| 1500 | 333298 | 15249 | 4806995 | 2 | 0 | 4725191 | 4725191 | 4725191 | 99.89 | 100 |

a. Two circular contigs indicates both the chromosome and the plasmid
b. NG50 - 50% of the entire assembly is contained in contigs or scaffolds equal to or larger than this value

**Table 3:** Assembly Data for MinION sequencing of *E. coli*

| Duration (min) | Reads | Subsampled Reads | Assembly Size | Circular Contigs<sup>a</sup> | Linear Contigs | Longest Contig | Longest Circular Contig | NG50<sup>b</sup> | Average identity in % |
| --- | --- | --- | --- | --- | --- | --- | --- | --- | --- |
| 15 | 8731 | 8731 | 1352560 | 0 | 19 | 154626 | 0 | 0 | 99.18 |
| 30 | 18053 | 18053 | 5141583 | 0 | 14 | 1565772 | 0 | 518218 | 99.63 |
| 60 | 35335 | 35335 | 5481126 | 1 | 0 | 5481126 | 5481126 | 5481126 | 99.82 |
| 120 | 62415 | 60362 | 5570410 | 1 | 1 | 5481662 | 5481662 | 5481662 | 99.87 |
| 240 | 103681 | 19589 | 5577045 | 2 | 0 | 5482542 | 5482542 | 5482542 | 99.89 |
| 480 | 164641 | 15265 | 5577346 | 2 | 0 | 5482831 | 5482831 | 5482831 | 99.90 |
| 960 | 317698 | 12941 | 5577818 | 2 | 0 | 5483284 | 5483284 | 5483284 | 99.91 |
| 1500 | 429909 | 12403 | 5577934 | 2 | 0 | 5483397 | 5483397 | 5483397 | 99.91 |

a. Two circular contigs indicates both the chromosome and the plasmid
b. NG50 - 50% of the entire assembly is contained in contigs or scaffolds equal to or larger than this value

The MinION sequencing data was assembled using a custom Nextflow^38^ workflow that utilized publicly-available tools. Fitlong length- and quality-based subsampling to a 75X target depth resulted in 28,492 reads for the *Salmonella* Bareilly isolate, which were assembled into two circular contigs, the chromosome and plasmid, with an average nucleotide identity of 99.87% and 100% coverage compared to the reference genome (Table 2). For the *E. coli* isolate, 19,589 subsampled reads produced two circular contigs, the chromosome and plasmid, with an average nucleotide identity of 99.89% compared to the available MiSeq data of the same bacterium (Table 3). The *Salmonella* Bareilly genome assembled into one chromosomal contig of 4,724,389 bp and one plasmid of 81,761 bp (Table 2). The *E.coli* O157:H7 genome assembled into one chromosomal contig of 5,482,542 bp and one plasmid of 94,503 bp (Table 3).

The final genome assemblies utilized two rounds of polishing using Nanopolish, which represented, by far, the most time-consuming and resource-intensive portion of the analysis workflow. However, it also increased the overall accuracy (Fig 1a) due to a decrease in both SNPs (Fig 1b) and chromosomal insertions or deletions (Fig 1c). The largest gains in accuracy were achieved from the first round of polishing, while much less but still noticeable improvement was achieved with the second round, particularly when examining completeness of genome annotation as measured by BUSCO. However, further rounds (>2) of polishing did not significantly impact the overall assembly (Fig 1). To demonstrate these data, relative time, central processing units (CPU), and memory consumption for each step of the workflow can be found in supplementary Table S1.

**Figure 1.**
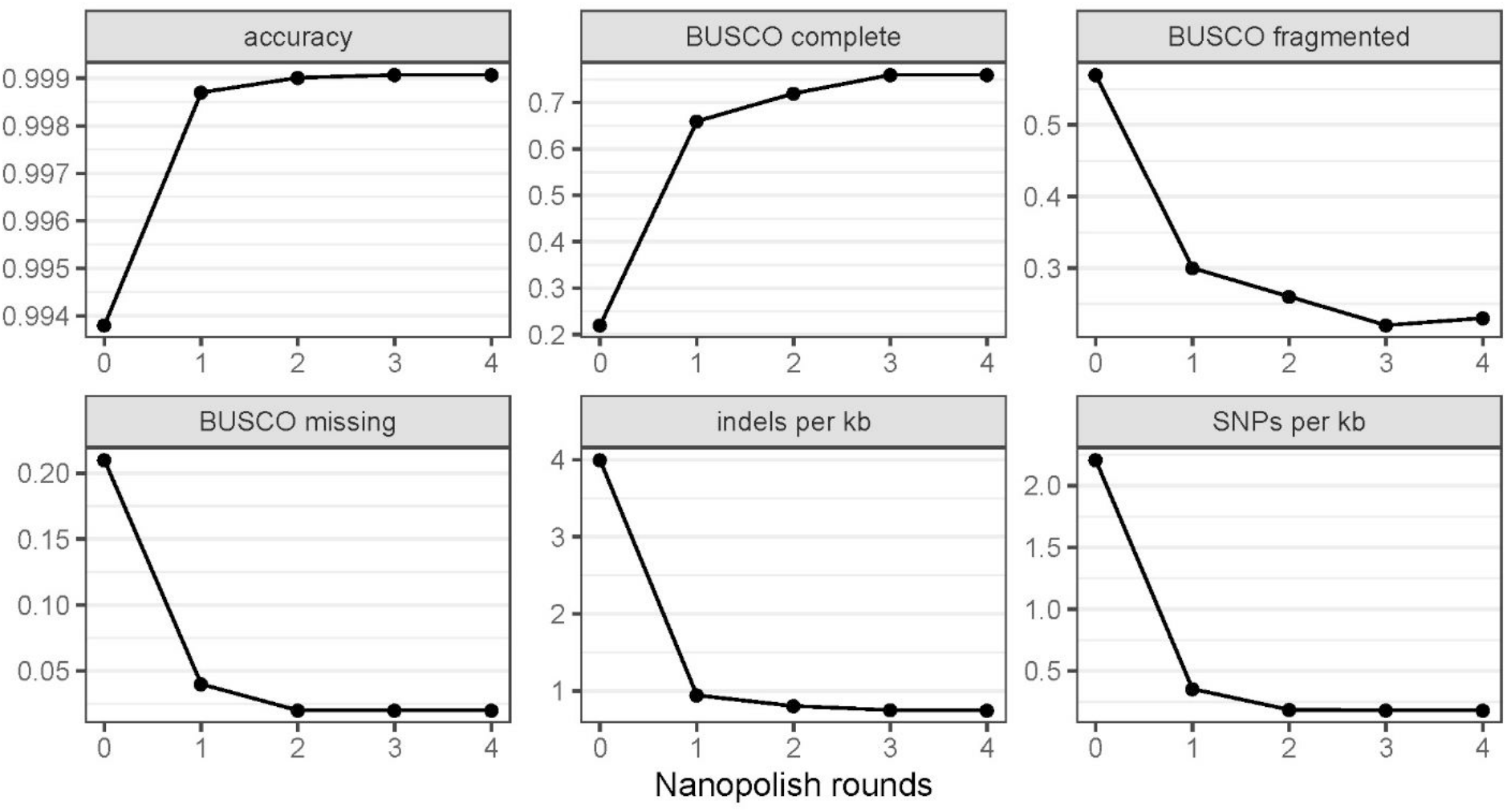
Polishing Results of the MinION-only Assemblies Using Multiple Rounds of Nanopolish. Due to the errors remaining in the MinION-only assemblies, a signal-level consensus software, Nanopolish, was used to increase the assembly accuracy. The overall accuracy, the Benchmarking Universal Single-Copy Orthologs (BUSCO) completeness, BUSCO Fragmented, BUSCO Missing, number of indels per kb, and number of SNPs per kb are shown after 0, 1, 2, 3 and 4 rounds of Nanopolish. After two rounds of polishing, the overall accuracy and the number of Indels and SNPs per kb did not considerably change.

For the *Salmonella* Bareilly assembly at 4 hours, the rate of single nucleotide polymorphisms (SNPs) per kilobase (kb) decreased from 2.41 to 0.42 after one round of polishing and to 0.26 after two rounds of polishing. At the same time point, the insertions or deletions (indels) per kb decreased from 3.91 to 1.14 after one round of polishing and to 1.03 after two rounds of polishing (Supplemental Table S2). For the *E. coli* assembly at the same time point, the SNPs per kb decreased from 2.20 to 0.37 after only one round of polishing and to 0.2 after two rounds of polishing. The indels per kb also decreased from 3.86 to 1 to 0.89 (Supplemental Table S3). Additionally, the BUSCO tool was used to further analyze the polished data to determine the completeness of the gene content based on quality and length of alignment. The “BUSCO completeness” (fraction of expected gene complement with full-length reading frames) value of both bacterial assemblies and the rounds of polishing were directly related, increasing from 21 and 23% for the *Salmonella* and *E. coli* assemblies, respectively, with no polishing to 65 and 69% after two rounds of polishing; the BUSCO fragmented (decreased length alignment of genes) and BUSCO missing (no significant matches) values decreased correspondingly (Supplemental Table S2 and S3).

### MinION Assembly Annotation

Both 4-hour, MinION-only assemblies, after two rounds of polishing with Nanopolish, were initially annotated using Geneious and the most closely related, published, annotated genomes for each bacterial species. The annotations were confirmed using the PGAP annotations of the final, corrected assemblies. Since the *Salmonella* Bareilly genome was already completed and closed, we confirmed that the genome annotation of the sequence produced by MinION was structurally identical to the annotation from the Illumina/PacBio sequence already published, for example, but not limited to, the two major serotyping antigens located on the chromosome: the flagellin FliC CDS and the O-antigen polymerase.

The presence of major virulence factors in the *E. coli* MinION-only assembly were identified, as well as genes that would cause possible antimicrobial resistance (Fig 2 and 3). The locus of enterocyte effacement (LEE), one of the major virulence factors of enterohemorrhagic *E. coli*^39,40^ that includes the gene intimin for adhesion and the type III secretion system, was annotated between positions 4,603,699 and 4,636,299 in this MinION-only assembly (Fig 2). Additionally, the genes expressing the Shiga toxins (Stx), responsible for causing host cell damage^39,41^, were annotated from position 3,181,004 to 3,181,963 for Stx subunit A and from position 3,180,723 to 3,180,992 for Stx2 subunit B (Fig 2). The multidrug resistance gene Mdf(A), which encodes a membrane protein that confers resistance to a multitude of clinically important drugs, including macrolides, lincosamides, and streptogramin B^42^, was also identified at position 1,012,477 to 1,013,709. No other genes or point mutations that would confer antimicrobial resistance were detected. Not only was the full-length chromosome of this *E.coli* O157:H7 isolate sequenced using MinION, but also the full-length pO157 (Fig 3). This plasmid also encodes *E. coli* O157-specific virulence factors^39^, such as hemolysin *(ehx)* identified at position 16,584 to 19,578, catalase-peroxidase (*katP*) at position 76,704 to 78,356, and the type II secretion system (T2SS) at position 64,056 to 85,694.

**Figure 2.**
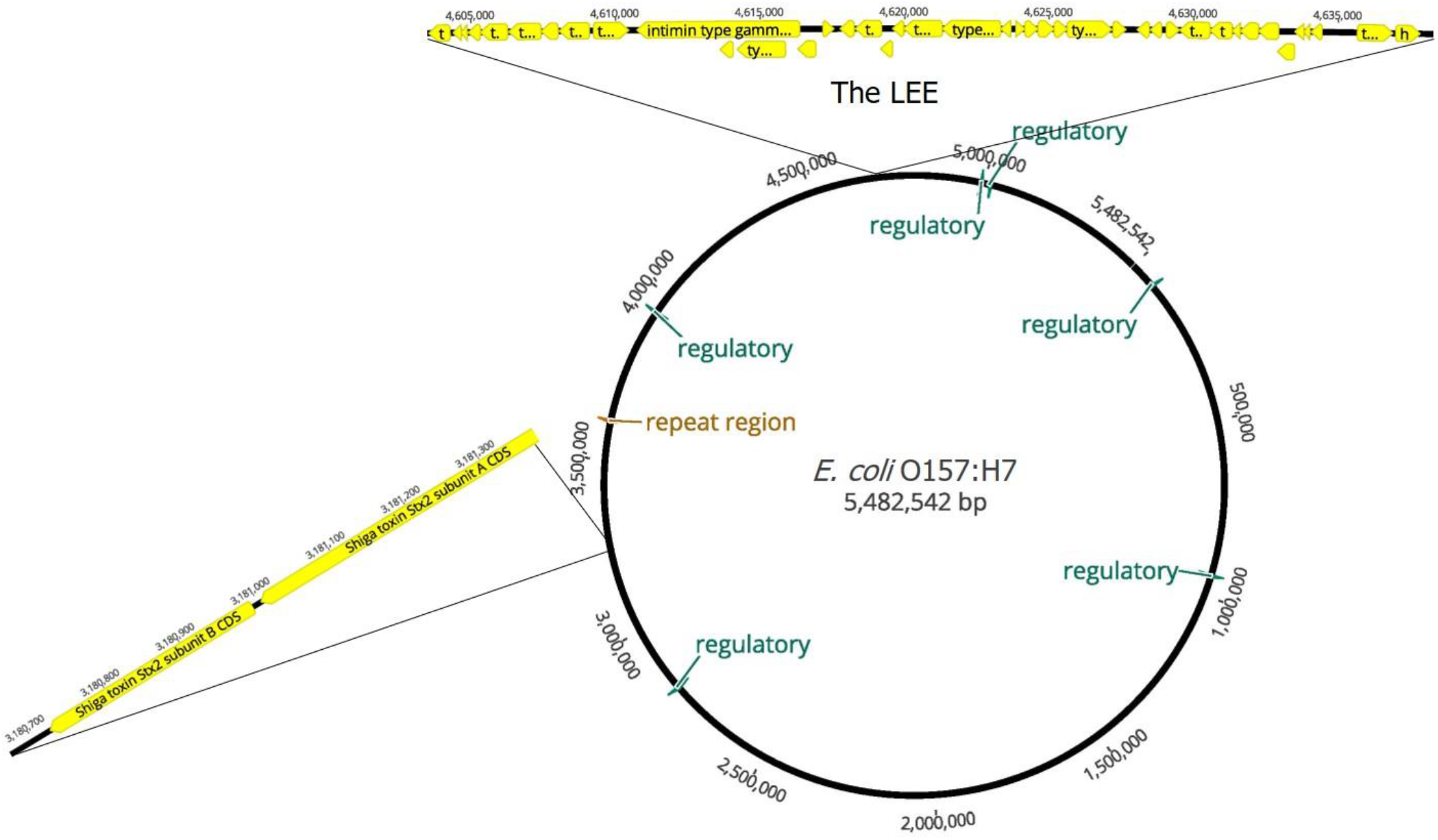
MinION Assembly of the *E. coli* chromosome. The *E.coli* O157:H7 chromosome was sequenced and assembled into a final consensus of 5,482,542 nucleotides. The annotation of the genome provided the location of 5,748 coding sequences (CDS), 106 tRNAs, 29 rRNAs, 6 regulatory regions, and 1 repeat regions. For imaging purposes, only the 6 regulatory regions (green), the one repeat region (brown) and the CDS of two virulence factors (yellow) are shown magnified. The LEE (locus of enterocyte effacement) is highlighted at position 4,603,699 to 4,636,299, and the Shiga Toxin subunits are shown at position 3,181,004 to 3,180,992.

**Figure 3.**
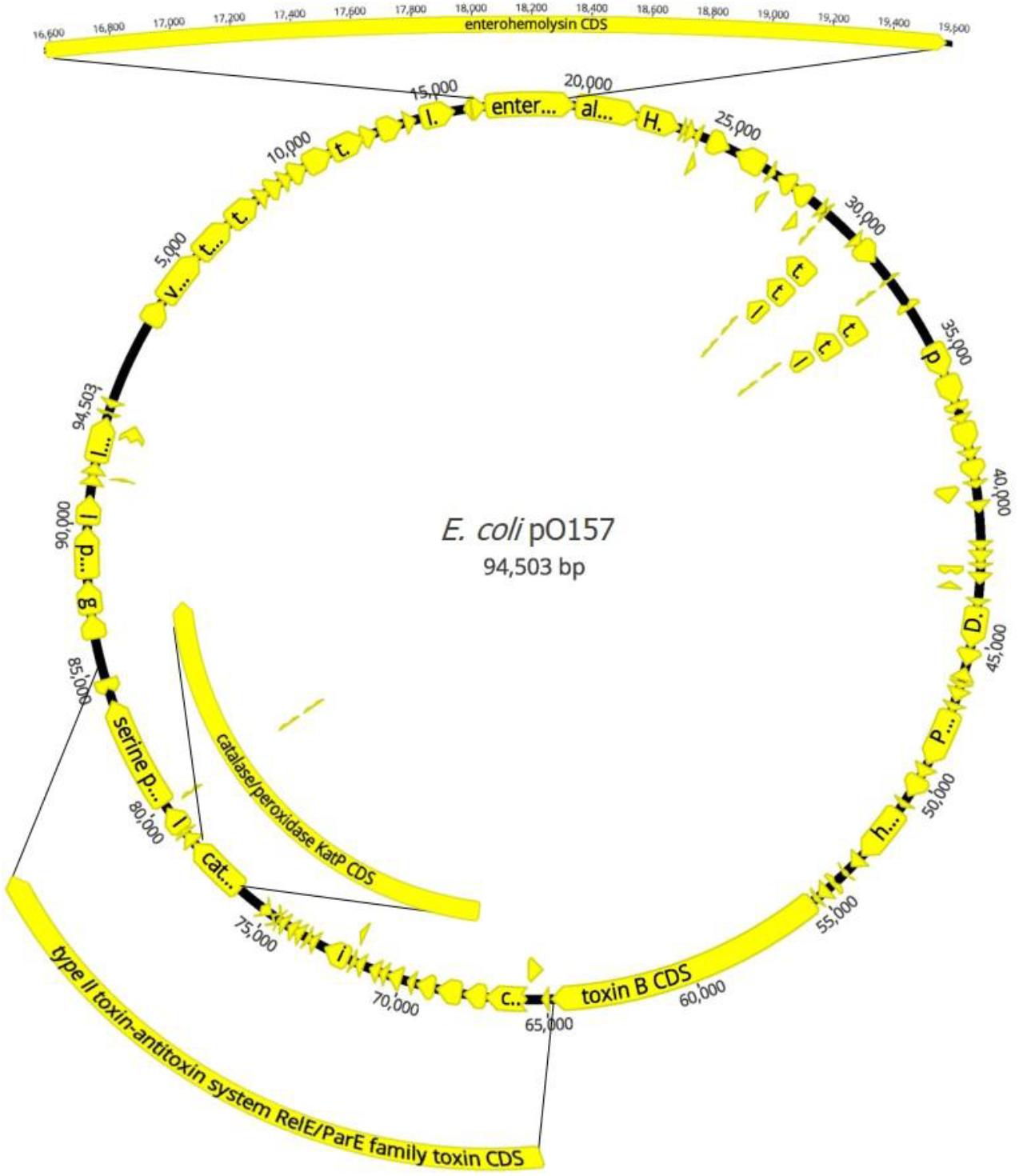
MinION Assembly of the *E. coli* pO157. The *E.coli* pO157 plasmid was sequenced and assembled into a final consensus of 94,503 nucleotides. The annotation shows all 124 coding sequences (CDS) in yellow. The CDS of three well-known virulence factors are highlighted: hemolysin (*ehx*) at position 16,584 to 19,578, catalase-peroxidase (*katP*) at position 76,704 to 78,356, and the type II secretion system (T2SS) at position 64,056 to 85,694.

### Additional Polishing of the MinION assemblies with MiSeq Data

This paper is primarily focused on optimizing sequencing run times and costs for food safety applications using nanopore-only approaches. However, for submission of final sequences to GenBank, the most accurate assemblies attainable were used. To this end, for both samples, assemblies produced using the full run length were utilized and further error-corrected using Pilon, together with available MiSeq data. Pilon utilizes the low error rate of Illumina reads mapped to the draft assembly to drastically improve the local accuracy of the final sequence. The error rate decreased for both samples after Pilon polishing, with accuracy rates of 99.99% and 100%, and BUSCO completeness rates of 99.7% and 99.99% for *Salmonella* and *E.coli*, respectively. There were also reductions in SNPs per kb to 0.002 and 0.001 and indels per kb to 0.008 and 0.002 for *Salmonella* and *E.coli*, respectively. The assembled, polished, and short-read error-corrected data from the full 25-hour run were the final assemblies annotated and submitted to GenBank (Accession numbers CP034177-CP034178 and CP035545-CP035546 with Bioproject PRJNA498670).

### Phylogenetic inference (SNP tree)

The constructed maximum likelihood SNPs trees are presented in Figure 4. The tree provided with the reference *Salmonella* datasets used for phylogenetic pipeline validation for foodborne pathogen surveillance^34^ is depicted in Figure 4A. To demonstrate the potential of the MinION-only sequencing for phylogenetic inference, the raw data for strain CFSAN000189 sequenced in this study was replaced with the data from our un-polished, MinION-only assemblies, and the tree was rebuilt from raw data using our pipeline (Figure 4B). For simplicity, only the 30 mins, 240 mins and 1500 mins timepoints were used for the reconstruction. The comparison between the trees built with the reference datasets and the tree utilizing the MinION-only data for the CSAFN000189 strain demonstrates complete topological congruence between both trees. The results using all eight time points showed identical topology except for the 15-minute time-point (data not shown).

**Figure 4.**
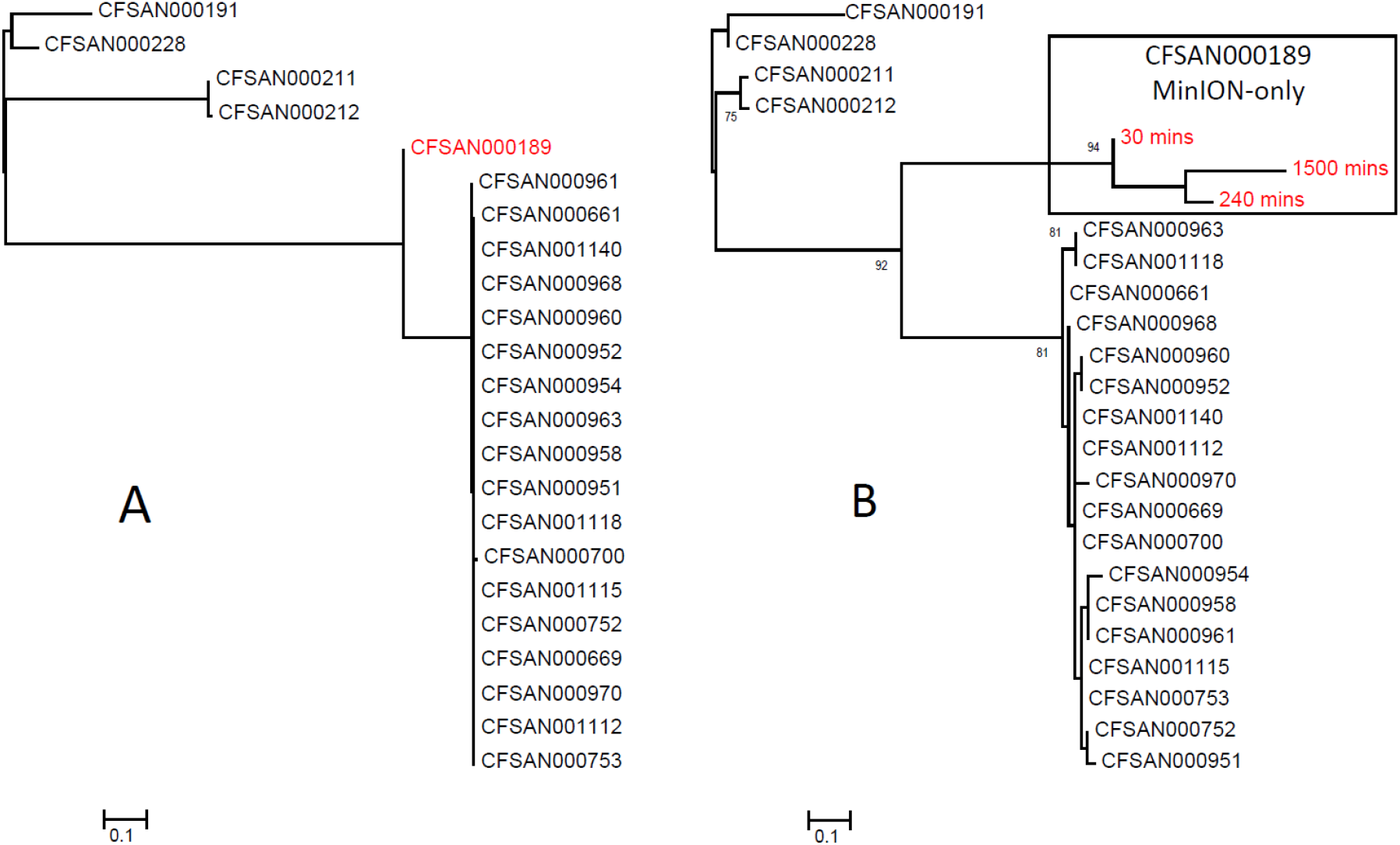
Maximum likelihood phylogenetic SNPs trees of Salmonella reference datasets. A) Maximum likelihood SNPs tree provided with the reference datasets from Timme et al., 2017 that was used for phylogenetic pipeline validation for foodborne pathogen surveillance ^34^. B) PhyML v3.0 was used to generate a maximum likelihood SNP tree for comparison purposes with twenty-two *Salmonella* reference datasets ^34^ and by replacing the CFSAN000189 data with SNPs from the 30 mins, 240 mins, and 1500 mins un-polished, MinION-only assemblies obtained in this study, which are enclosed in a rectangle and are shown in red. The HYK85 model was used with 500 bootstraps, and branches with less than 60% support were collapsed. There were a total of 64 positions in the final dataset. The tree is drawn to scale, with branch lengths measured in the number of substitutions per site.

## Discussion

In this study, we demonstrate that long-read, nanopore sequencing technology can be used as a single tool to sequence full length bacterial chromosomes and plasmids. Utilizing publicly-available tools in a customized workflow, the MinION-only data produced assembled sequences with as little as 0.1% error rate, which is 0.4% and 3.1% lower than previous reports^9,19^. The tools used in our customized bioinformatic workflow are publicly-available^24,25^. The workflow was optimized and tailored to the specific data being analyzed, and the additionally used local code is also made available on Github.

Using MinION sequencing alone, two completely closed contigs, one chromosome and one plasmid, for each pathogen were assembled. This capability and the low cost make the MinION highly accessible as both a primary sequencing platform, as well as a secondary platform to complement laboratories’ existing sequencing infrastructure. The initial investment required for the MinION is drastically lower than other sequencing technologies (currently, a starter pack is $1000, which includes the instrument, two flow cells, usually $500 each, and the respective wash, sequencing, and library loading kit). Additionally, each flow cell can be used for multiple runs, and samples can be multiplexed together per run to reduce the cost^20,43^. Based on the results of barcoding and simultaneous sequencing of two whole bacterial genomes and plasmids shown here, we estimate that six bacterial samples could be multiplexed together to further decrease cost and sequenced in approximately 16 hours to obtain complete genomic data with high accuracy.

The effects of increased sequencing run lengths, different criteria and weights to subsample data for assembly, and increased rounds of polishing were examined for their effect on the final assembly completeness and accuracy. It was observed that the nanopore reads were long enough on average, and that over-aggressive length-based filtering resulted in reduced representation. Such extensive subsampling would result in less complete assembly of small plasmids, which can contain virulence factors of great interest for diagnostic and food safety purposes. It therefore proved critical to evaluate filtering and subsampling criteria to take full advantage of the technology.

Our results suggest that at least one round of polishing with Nanopolish is needed to achieve acceptable accuracy, and a second round provides additional improvement if the near-doubling of the analysis time is warranted. The data in Table S1 is provided when only one core is utilized, but due to the availability of high-performance computers, the analysis time for two rounds of polishing can decrease to 6 hours using 124 cores, for example. In MinION-only assemblies, it is known that putative pseudogenes caused by systematic indel errors (often near homopolymeric tracts^19,44^), leading to reading frame shifts can be an issue, as evident from the “BUSCO fragmented” column in Supplementary tables S2 and S3. Even after polishing, this value was observed to be greater than 20% of expected coding genes, which must be taken into consideration during annotation. However, the polished assemblies, with only 0.1% error, still reveal serotype and important genes responsible for the virulence, metabolism, defense, and pathogenesis of the bacterium.

In outbreak situations, a rapid turn-around time is necessary. Therefore, polymerase chain reaction (PCR), real-time PCR assays, and other rapid diagnostic assays are still employed. However, WGS is far more powerful and informative and has become routine in use as it can be coupled with proper bioinformatics analysis to provide complete genome sequences in a couple of days^2^. With the MinION platform and sufficient computational resources (which can be cloud-based and thus widely available), basecalled sequence data can potentially be analyzed in near-real-time as it comes off the machine^45^. Therefore, the MinION can be used for rapid diagnostics as initial sequencing data from pure cultures can be provided in approximately 9 to 10 hours^46^. The complete MinION data can be further analyzed and polished after the entire sequencing run to obtain accurate, whole genomes that provide detailed data on subtyping, virulence genes, antimicrobial resistance genes, and other genetic characteristics. Same-day detection of antimicrobial resistance genes with 99.75% accuracy (with polishing) after enriching for plasmid DNA and MinION sequencing has been recently demonstrated^47^. Here, we also demonstrate the potential of the un-polished, MinION-only results for rapid phylogenetic inference with identical topology as compared to the reference published dataset after only 30 mins of sequencing, with consistent results at all other analyzed time points. The application of the technology for epidemiological tracing during real field outbreak remains to be tested and verified utilizing more samples.

In conclusion, this low-cost, rapid, random-priming nanopore sequencing approach, coupled with our customized workflow, provides sufficient data where complete genomes, including plasmids, can be assembled into a single contiguous sequence with 99.89% accuracy. These data provided both gene identification and genomic organization without the need for additional sequencing tools to close gaps that are required by other sequencing methods. We were able to successfully sequence complete bacterial genomes with the lowest error rate reported-to-date using a single sequencing method and to demonstrate the potential of these results for epidemiological inference and outbreak tracing. As the nanopore chemistry and bioinformatics continue to evolve, this method is promising in providing a sufficient amount of accurate data to complement the current sequencing methods by resolving repetitive regions of the genome, which will be instrumental in increasing the number of available complete genome assemblies.

## Acknowledgments

The authors would like to thank Drs. Ruth Timme and James Pettengill with the FDA Center for Food Safety and Applied Nutrition for the *Salmonella* Bareilly isolate used in this manuscript. This work was supported by U.S. Department of Agriculture, ARS CRIS Project 6612-32000-072.

The mention of trade names or commercial products in this publication is solely for the purpose of providing specific information and does not imply recommendation or endorsement by the U.S. Department of Agriculture. The USDA is an equal opportunity provider and employer.

## Author Contributions

Conceptualization – T. Taylor, E. DeJesus, G. Tillman, C. Afonso.; Methodology - T. Taylor and E. DeJesus.; Software - J. Volkening and M. Simmons.; Formal Analysis - T. Taylor, J. Volkening, M. Simmons, K. Dimitrov; Resources – G. Tillman, D. Suarez, C. Afonso.; Original Draft Preparation - T. Taylor; Review & Editing - T. Taylor, J. Volkening, E. DeJesus, M. Simmons, K. Dimitrov, G. Tillman, D. Suarez, C. Afonso; Supervision, Project Administration, Funding Acquisition - D. Suarez, C. Afonso.

## Data availability

The final assemblies generated during the current study are available in GenBank (Accession CP034177-CP034178 and CP035545-CP035546). The raw data generated during the current study are available from the corresponding author on reasonable request.

