## Supplemental Data for "Rapid, multiplexed, whole genome and plasmid sequencing of foodborne pathogens using long-read nanopore technology"

**Supplemental Figure 1. MinION sequencing run time versus accuracy of polished (2x) assemblies.** The raw MinION sequencing data for both isolates were subsampled based on cumulative run time in order to simulate the effect of run length on polished assembly quality. At the 240-minute time point, two circularized contigs, one chromosome and one plasmid, from both isolates were fully sequenced, and the accuracy, after two rounds of polishing, levels off and does not significantly increase with longer run times.

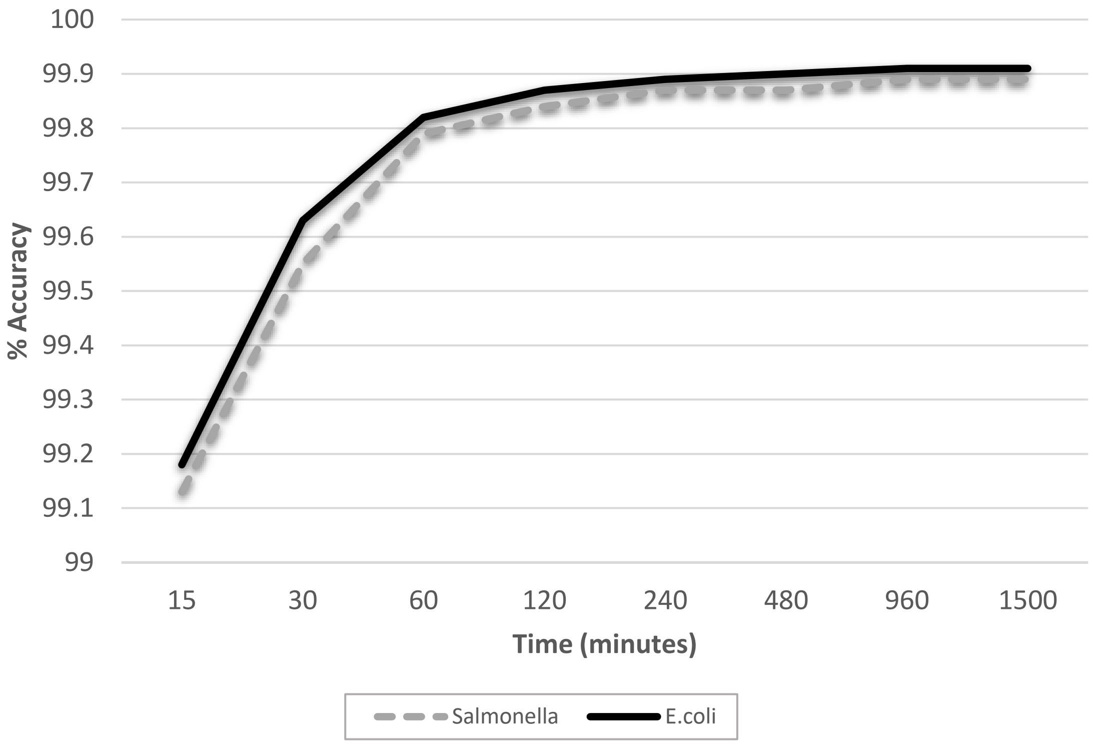

**Supplemental Table S1: Resource usage for main steps in the data analysis.** The amount of central processing unit hours and the amount of memory used in Gigabytes (GB) for steps of the workflow is provided for assembly without polishing and assembly after two rounds of polishing.

| **Processor usage (CPU hours)** | | | |
|---|---|---|---|
| **Step** | **No Nanopolish** | **One Round of Nanopolish** | **Two rounds of Nanopolish** |
| Unicycler | 8.8 | 8.8 | 8.8 |
| Nanopolish | 0 | 359.6 | 729.7 |
| Other | 4.5 | 4.5 | 4.5 |
| **Maximum memory usage (GB)** | | | |
| **Step** | **No Nanopolish** | **One round of Nanopolish** | **Two Rounds of Nanopolish** |
| Unicycler | 5.4 | 5.4 | 5.4 |
| Nanopolish | 0 | 1 | 1 |
| Other | 5.2 | 5.2 | 5.2 |

**Supplemental Table S2:** ***Salmonella*** **Bareilly MinION sequencing data analyzed for completeness and accuracy before and after two rounds of polishing.** After two rounds of polishing, the average identity at each sequencing duration time-point increases, while the SNPs and Indels per kb decrease. The percentage of BUSCO complete also increases after two rounds of polishing, while the BUSCO fragmented and missing portions of the genome decrease. The 240-minute sequencing duration is highlighted, since all presented analyses in the text are based on the data collected at this time-point.

| Seq Duration (min) | Reference Coverage | Avg. ID | rel[a] | t[b] | inv[c] | ins[d] | ins sum | SNPs/ kb[e] | Indels/ kb[f] | BUSCO complete[g,h] | BUSCO fragmented[g,i] | BUSCO missing[g,j] |
|---|---|---|---|---|---|---|---|---|---|---|---|---|
| **No Polishing** | | | | | | | | | | | | |
| **15** | 24.36 | 98.6 | 0 | 0 | 1 | 12 | 506 | 4.04 | 9.43 | 0.02 | 0.07 | 0.91 |
| **30** | 95.38 | 99.05 | 0 | 0 | 1 | 10 | 292 | 3.01 | 6.43 | 0.14 | 0.51 | 0.35 |
| **60** | 98.4 | 99.3 | 0 | 0 | 1 | 0 | 0 | 2.54 | 4.46 | 0.19 | 0.59 | 0.22 |
| **120** | 100 | 99.32 | 0 | 0 | 1 | 1 | 3613 | 2.43 | 4.39 | 0.2 | 0.57 | 0.22 |
| **240** | 100 | 99.37 | 0 | 0 | 1 | 1 | 3612 | 2.41 | 3.91 | 0.21 | 0.57 | 0.22 |
| **480** | 100 | 99.37 | 0 | 0 | 1 | 1 | 3618 | 2.42 | 3.89 | 0.21 | 0.55 | 0.24 |
| **960** | 100 | 99.36 | 0 | 0 | 1 | 1 | 3612 | 2.38 | 4.05 | 0.2 | 0.57 | 0.23 |
| **1500** | 100 | 99.37 | 0 | 0 | 1 | 2 | 3606 | 2.4 | 3.85 | 0.23 | 0.55 | 0.22 |
| **One Round of Polishing** | | | | | | | | | | | | |
| **15** | 24.36 | 99.1 | 0 | 0 | 1 | 11 | 494 | 2.19 | 6.52 | 0.04 | 0.11 | 0.85 |
| **30** | 95.38 | 99.52 | 0 | 0 | 1 | 10 | 292 | 1.22 | 3.48 | 0.32 | 0.5 | 0.18 |
| **60** | 98.4 | 99.77 | 0 | 0 | 1 | 0 | 0 | 0.6 | 1.72 | 0.46 | 0.44 | 0.1 |
| **120** | 100 | 99.81 | 0 | 0 | 1 | 1 | 3610 | 0.49 | 1.38 | 0.54 | 0.38 | 0.08 |
| **240** | 100 | 99.84 | 0 | 0 | 1 | 1 | 3616 | 0.42 | 1.14 | 0.61 | 0.32 | 0.07 |
| **480** | 100 | 99.86 | 0 | 0 | 1 | 1 | 3616 | 0.4 | 1.08 | 0.61 | 0.33 | 0.06 |
| **960** | 100 | 99.85 | 0 | 0 | 1 | 1 | 3612 | 0.44 | 1.02 | 0.62 | 0.33 | 0.06 |
| **1500** | 100 | 99.86 | 0 | 0 | 1 | 2 | 3610 | 0.41 | 1.01 | 0.62 | 0.31 | 0.06 |
| **Two Rounds of Polishing** | | | | | | | | | | | | |
| **15** | 24.36 | 99.13 | 0 | 0 | 1 | 11 | 492 | 2.06 | 6.35 | 0.05 | 0.12 | 0.84 |
| **30** | 95.38 | 99.55 | 0 | 0 | 1 | 10 | 292 | 1.1 | 3.33 | 0.34 | 0.48 | 0.18 |
| **60** | 98.4 | 99.79 | 0 | 0 | 1 | 0 | 0 | 0.48 | 1.61 | 0.5 | 0.42 | 0.08 |
| **120** | 100 | 99.84 | 0 | 0 | 1 | 1 | 3610 | 0.34 | 1.26 | 0.58 | 0.35 | 0.07 |
| **240** | 100 | 99.87 | 0 | 0 | 1 | 1 | 3616 | 0.26 | 1.03 | 0.65 | 0.3 | 0.05 |
| **480** | 100 | 99.87 | 0 | 0 | 1 | 1 | 3616 | 0.24 | 0.99 | 0.66 | 0.29 | 0.05 |
| **960** | 100 | 99.89 | 0 | 0 | 1 | 1 | 3612 | 0.23 | 0.89 | 0.67 | 0.28 | 0.04 |
| **1500** | 100 | 99.89 | 0 | 0 | 1 | 2 | 3610 | 0.23 | 0.86 | 0.69 | 0.27 | 0.04 |

a. Relocations – rearrangement of genetic material within a chromosome or between chromosomes
b. Translocations- rearrangement of parts between nonhomologous chromosomes
c. Inversions - rearrangement in which a segment of a chromosome is reversed end to end
d. Insertions - the addition of a larger nucleotide sequence into a chromosome
e. SNPs/kb – single nucleotide polymorphisms per kilobase
f. Indels/kb – insertions or deletions per kilobase
g. BUSCO- Benchmarking Universal Single-Copy Orthologs
h. Complete-fraction of expected gene complement with full-length reading frames
i. Fragmented- decreased length alignment of genes
j. Missing- no significant matches

**Supplemental Table S3: *E. coli* MinION sequencing data analyzed for completeness and accuracy before and after two rounds of polishing.** After two rounds of polishing, the average identity at each sequencing duration time-point increases, while the SNPs and Indels per kb decrease. The percentage of BUSCO complete also increases after two rounds of polishing, while the BUSCO fragmented and missing portions of the genome decrease. The 240-minute sequencing duration is highlighted, since all presented analyses in the text are based on the data collected at this time-point.

| Seq Duration (min) | Avg ID | SNPs/ kb[a] | indels/ kb[b] | BUSCO complete[c,d] | BUSCO fragmented[c,e] | BUSCO missing[c,f] |
|---|---|---|---|---|---|---|
| **No polishing** | | | | | | |
| **15** | 98.62 | 4.06 | 9.69 | 0.01 | 0.06 | 0.93 |
| **30** | 99.16 | 2.67 | 5.71 | 0.13 | 0.5 | 0.37 |
| **60** | 99.36 | 2.31 | 4.07 | 0.2 | 0.57 | 0.23 |
| **120** | 99.4 | 2.22 | 3.74 | 0.23 | 0.55 | 0.22 |
| **240** | 99.39 | 2.22 | 3.86 | 0.23 | 0.54 | 0.23 |
| **480** | 99.38 | 2.21 | 4 | 0.22 | 0.57 | 0.21 |
| **960** | 99.41 | 2.25 | 3.71 | 0.23 | 0.55 | 0.22 |
| **1500** | 99.4 | 2.24 | 3.72 | 0.22 | 0.58 | 0.2 |
| **One round of Nanopolish** | | | | | | |
| **15** | 99.13 | 2.11 | 6.61 | 0.04 | 0.11 | 0.86 |
| **30** | 99.6 | 1.02 | 2.96 | 0.35 | 0.46 | 0.19 |
| **60** | 99.79 | 0.55 | 1.51 | 0.51 | 0.41 | 0.08 |
| **120** | 99.85 | 0.39 | 1.14 | 0.58 | 0.35 | 0.07 |
| **240** | 99.86 | 0.37 | 1 | 0.64 | 0.31 | 0.04 |
| **480** | 99.87 | 0.35 | 0.94 | 0.66 | 0.3 | 0.04 |
| **960** | 99.88 | 0.35 | 0.87 | 0.66 | 0.3 | 0.04 |
| **1500** | 99.88 | 0.35 | 0.87 | 0.64 | 0.31 | 0.05 |
| **Two rounds of Nanopolish** | | | | | | |
| **15** | 99.18 | 1.92 | 6.31 | 0.04 | 0.11 | 0.86 |
| **30** | 99.63 | 0.88 | 2.81 | 0.37 | 0.44 | 0.19 |
| **60** | 99.82 | 0.41 | 1.39 | 0.57 | 0.36 | 0.06 |
| **120** | 99.87 | 0.26 | 1.04 | 0.64 | 0.3 | 0.06 |
| **240** | 99.89 | 0.2 | 0.89 | 0.69 | 0.28 | 0.04 |
| **480** | 99.9 | 0.19 | 0.8 | 0.72 | 0.26 | 0.02 |
| **960** | 99.91 | 0.19 | 0.75 | 0.73 | 0.25 | 0.02 |
| **1500** | 99.91 | 0.18 | 0.74 | 0.73 | 0.24 | 0.03 |

a. SNPs/kb – single nucleotide polymorphisms per kilobase
b. Indels/kb – insertions or deletions per kilobase
c. BUSCO- Benchmarking Universal Single-Copy Orthologs
d. Complete-fraction of expected gene complement with full-length reading frames
e. Fragmented- decreased length alignment of genes
f. Missing- no significant matches

**Supplemental Table S4: Salmonella outbreak isolates used as reference data set for phylogenetic analysis.**

| Strain | GenBank Assembly | SRA Accession | Outbreak year-Location |
|---|---|---|---|
| CFSAN000212 | GCA_000748245.1 | SRR500494 | 2005-UAE |
| CFSAN000211 | GCA_000698715.1 | SRR498373 | 2005-Thailand |
| CFSAN000191 | GCA_000698635.1 | SRR498369 | 2005-Thailand |
| CFSAN000228 | GCA_000748565.1 | SRR500493 | 2006-Taiwan |
| CFSAN000189 | GCA_000439415.1 | SRR498276 | 2003-India |
| CFSAN000963 | GCA_000749295.1 | SRR498436 | 2012-USA |
| CFSAN001118 | GCA_000748065.1 | SRR1258443 | 2012-Indonesia |
| CFSAN000661 | GCA_000698515.1 | SRR498397 | 2012-USA |
| CFSAN000968 | GCA_000698535.1 | SRR498442 | 2012-USA |
| CFSAN001140 | GCA_000748085.1 | SRR1258440 | 2012-India |
| CFSAN001112 | GCA_000748025.1 | SRR1258439 | 2012-India |
| CFSAN000970 | GCA_000749415.1 | SRR498444 | 2012-USA |
| CFSAN000669 | GCA_000749005.1 | SRR498399 | 2012-USA |
| CFSAN000700 | GCA_000749045.1 | SRR498402 | 2012-USA |
| CFSAN000752 | GCA_000749065.1 | SRR498403 | 2012-USA |
| CFSAN000951 | GCA_000749145.1 | SRR498422 | 2012-USA |
| CFSAN000753 | GCA_000749085.1 | SRR498404 | 2012-USA |
| CFSAN001115 | GCA_000748045.1 | SRR1258442 | 2012-India |
| CFSAN000954 | GCA_000748405.1 | SRR498425 | 2012-USA |
| CFSAN000952 | GCA_000749165.1 | SRR498423 | 2012-USA |
| CFSAN000960 | GCA_000748505.1 | SRR498433 | 2012-USA |
| CFSAN000958 | GCA_000748465.1 | SRR498431 | 2012-USA |
| CFSAN000961 | GCA_000748525.1 | SRR498434 | 2012-USA |
